# Transcriptional heterogeneity in human diabetic foot wounds

**DOI:** 10.1101/2023.02.16.528839

**Authors:** Teresa Sandoval-Schaefer, Quan Phan, Biraja C. Dash, Alexandre J. Prassinos, Kaiti Duan, Michael I. Gazes, Steven D. Vyce, Ryan Driskell, Henry C. Hsia, Valerie Horsley

## Abstract

Wound repair requires the coordination of multiple cell types including immune cells and tissue resident cells to coordinate healing and return of tissue function. Diabetic foot ulceration is a type of chronic wound that impacts over 4 million patients in the US and over 7 million worldwide (Edmonds et al., 2021). Yet, the cellular and molecular mechanisms that go awry in these wounds are not fully understood. Here, by profiling chronic foot ulcers from non-diabetic (NDFUs) and diabetic (DFUs) patients using single-cell RNA sequencing, we find that DFUs display transcription changes that implicate reduced keratinocyte differentiation, altered fibroblast function and lineages, and defects in macrophage metabolism, inflammation, and ECM production compared to NDFUs. Furthermore, analysis of cellular interactions reveals major alterations in several signaling pathways that are altered in DFUs. These data provide a view of the mechanisms by which diabetes alters healing of foot ulcers and may provide therapeutic avenues for DFU treatments.

## Introduction

Tissue repair requires the coordination of multiple cell types to restore the cellular structures that allow a tissue to maintain its function (Eming et al., 2014). After an injury, an inflammatory response occurs in mammalian tissues that involves blood-derived neutrophil and monocyte-derived inflammatory macrophages that clear pathogens and cellular debris. Eventually, neutrophils and inflammatory macrophages decline, and the proliferative phase of tissue repair ensues as additional waves of monocytes are recruited to wounds and differentiate into antiinflammatory macrophages. In the skin, anti-inflammatory macrophages coordinate tissue-resident keratinocytes, fibroblasts, and blood vessels to repair the damaged epidermis and underlying dermis. Finally, the repaired skin is remodeled as cells are pruned and form a scar.

Diabetes leads to defects in multiple cell types and pathologies including neuropathy and vasculopathy (Brem and Tomic-Canic, 2007). Wound repair in diabetic mouse models and studies of human diabetic foot ulcers (DFUs) have revealed that diabetic wounds display persistent inflammatory macrophages and defective keratinocyte migration, angiogenesis, and extracellular matrix protein deposition (Joshi et al., 2020, Mirza and Koh, 2011, Shook et al., 2016). Diabetic foot ulceration impacts 15% of patients with diabetes, impairing their quality of life and resulting in lower extremity amputations (Edmonds et al., 2021). With the increase in diabetes worldwide, therapies that promote healing in DFUs are a major clinical need. These therapies would reduce the burden of DFU care on the healthcare system and patient morbidity and mortality, which occurs within 5 years for 50% of the patients with DFUs (Edmonds et al., 2021).

A major barrier in developing effective clinical therapies is the incomplete characterization of the cells and molecules that occur in wounds of volar skin, such as the foot. Volar (palmoplantar) skin has a unique structure compared to non-volar, dorsal skin: it contains more epidermal layers and different keratin expression (Swensson et al., 1998), lacks hair follicles, and has reduced pigmentation (Tsai et al., 2022). Due in part to the lack of large areas of volar skin in mammalian model organisms, our understanding of wound repair mechanisms in volar skin and how diabetes impedes the healing of this tissue is limited.

Here, we utilize single-cell RNA sequencing (scRNASeq) to define cellular and molecular heterogeneity of foot wounds in non-diabetic and diabetic patients. We provide a comprehensive atlas of how transcriptional changes in keratinocytes, fibroblasts, immune cells, and endothelial cells are altered in DFUs compared to foot wounds from non-diabetic individuals. We also identify alterations in cellular communication that characterize healing of diabetic foot wounds, which highlights defects in several specific signaling pathways. Together, these data provide several potential therapeutic avenues to promote healing in DFUs. Furthermore, we share these data through our easily accessible webtools: https://skinregeneration.org/data/.

## Results

### Identification of cellular heterogeneity in diabetic foot wounds

To understand the cellular and transcriptional heterogeneity associated with diabetic foot wound repair, we utilized scRNASeq of cells isolated from foot wounds of a single non-diabetic individual at different times and 5 diabetic patients. After mechanical and enzymatic dissociation, filtration, and removal of red blood cells, single-cell suspensions were submitted through a pipeline for scRNAseq using the 10X Genomics platform (**Figure 1A**).

**Figure 1.**
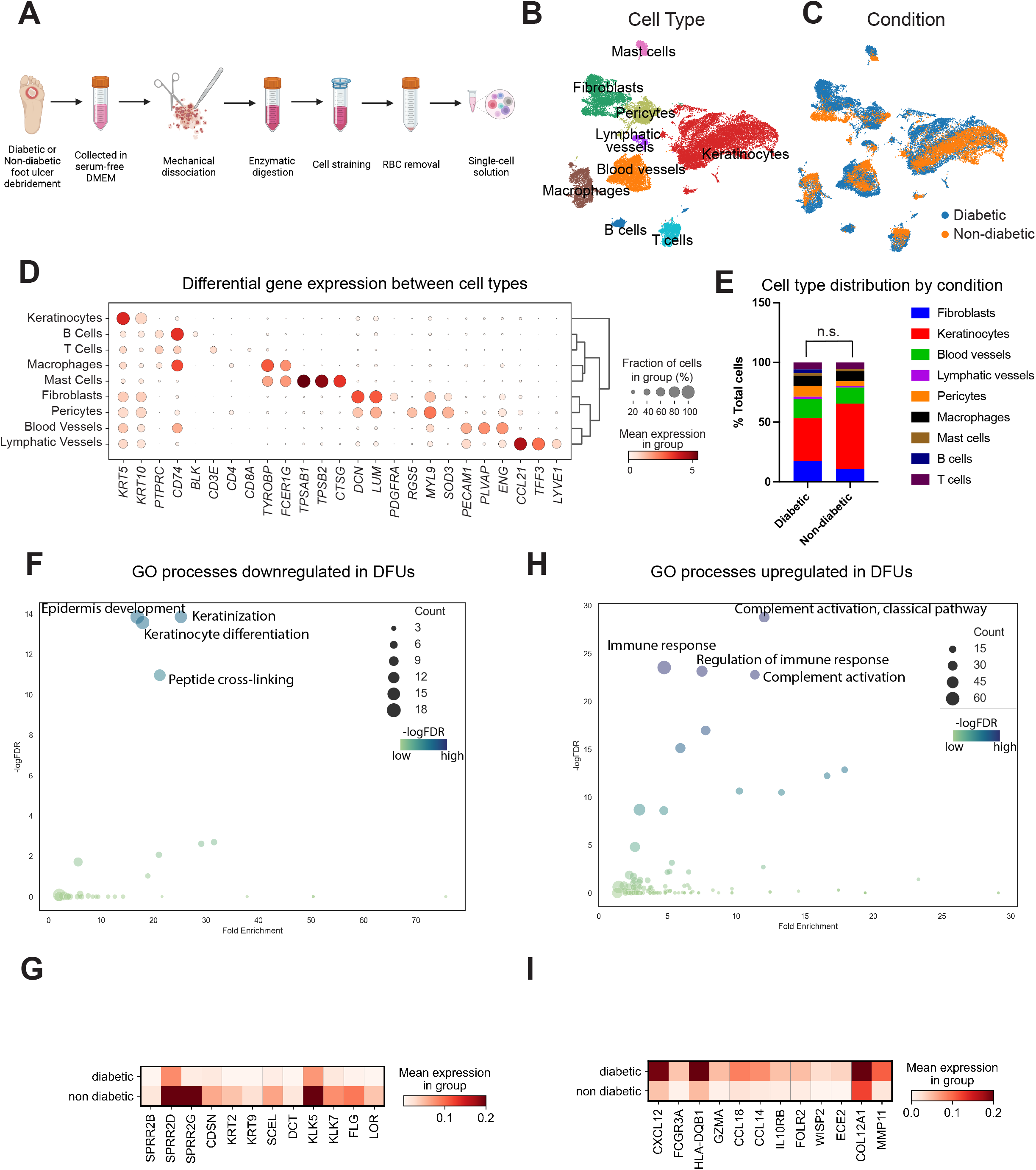
Single-cell RNA sequencing reveals cellular heterogeneity and differential gene expression between DFU and NDFU. **(A)** Schematic representation of the experimental procedure. DFU and NDFU specimens were harvested and processed separately following the same procedure. **(B)** Unbiased UMAP clustering of 27998 cells reveals 9 cellular clusters. Clusters are identified by color and cell type. **(C)** Stacked bar graph showing the percentage of each cell type by condition. (n.s., not significant) **(D)** Dot plot showing the differential gene expression of markers used to identify each cell cluster. The area of the circles indicates the proportion of cells expressing the gene, and the color intensity reflects the expression intensity. **(E)** UMAP clustering of all cells by condition showing the distribution of DFU and NDFU cells in clusters. **(F)** GO analysis of biological processes downregulated in DFU by -logFDR and Fold Enrichment. The area of the circles indicates the gene count, and the color intensity reflects the -logFDR power. **(G)** Matrix plot showing the differential expression of selected genes that are downregulated in DFU. The color intensity reflects the expression intensity. **(H)** GO analysis of biological processes upregulated in DFU by -logFDR and Fold Enrichment. The area of the circles indicates the gene count, and the color intensity reflects the -logFDR power. **(I)** Matrix plot showing the differential expression of selected genes that are upregulated in DFU. The color intensity reflects the expression intensity. DFU, diabetic foot ulcers. FDR, false discovery rate. NDFU, non-diabetic foot ulcers. UMAP, uniform manifold approximation and projection. Panel (A) was created with BioRender.com

Our initial analysis provided an overview of the diverse foot skin populations by integrating all 8 samples together from diabetic and nondiabetic individuals. Data analysis of 27,998 cells from all 8 samples were integrated using Scanorama batch correction. The transcriptomes of these cells were visualized on 2D graphs using dimension reduction technique Uniform Manifold Approximation and Projection (UMAP) (Becht et al., 2018). Clusters of unique cell types were identified using community detection algorithm *leiden* and differential gene expression profiles, which were then assigned 9 clusters with distinct gene expression profiles (**Figure 1B**). Comparing known genes with the most representative expressed mRNAs of each cluster (**Figures 1D**, **S1B**) revealed the identity of the cell clusters. We identified keratinocytes (*K5+, K10+*), several fibroblast populations (*DCN+, LUM+, PDGFRA+/-*), endothelial (blood (*PECAM+, PLVAP+*) and lymphatic (*CCL21+, LYVE1+*)) cells, and immune cells including macrophages (*TYROBP+*), mast cells (*TPSAB1+, TPSB2+, CTSG+*), T (*CD3E+, CD8A+*) and B (*CD74+, BLK+*) cells (**Figures 1B, 1D, S1B**). All cell clusters contained cells from all donors (**Figure S1A**) except B cells, which were only prevalent in one DFU sample and thus were removed from further analysis (Figure S1A, 4C). Furthermore, we found that the cell numbers were inconsistent between samples, which is consistent with the variability of human samples and library processing of scRNAseq data (**Figure 1C, Table 2**)(Ziegenhain et al., 2018).

UMAP visualization of diabetic vs non-diabetic cells revealed differences in gene expression in major clusters (**Figure 1E**). To assess these differences, we analyzed gene expression changes in diabetic samples compared to non-diabetic samples. We found that epidermal development and differentiation were significantly downregulated in the diabetic foot ulcer cells including genes involved in response to wounding (*K17, CASP14*) and genes involved in keratinocyte differentiation (*SPRR* genes, *TGM5*) (**Figure 1F, 1G, Table 3**). Interestingly, genes involved in inflammation and ECM proteolysis were significantly upregulated in diabetic samples (**Figure 1H, 1I, Table 3**).

### Keratinocyte adhesion, differentiation, and ECM regulation are transcriptionally altered in diabetic foot wounds

To explore differences in keratinocytes in diabetic and nondiabetic foot wounds in more detail, we clustered keratinocytes independently from other cell types (**Figure 2A-2E**). We noted that diabetic wounds displayed distinct clustering in several clusters including a cluster expressing migration genes, a basal cell population, and several differentiating keratinocyte clusters (**Figure 2A, 2B**).

**Figure 2.**
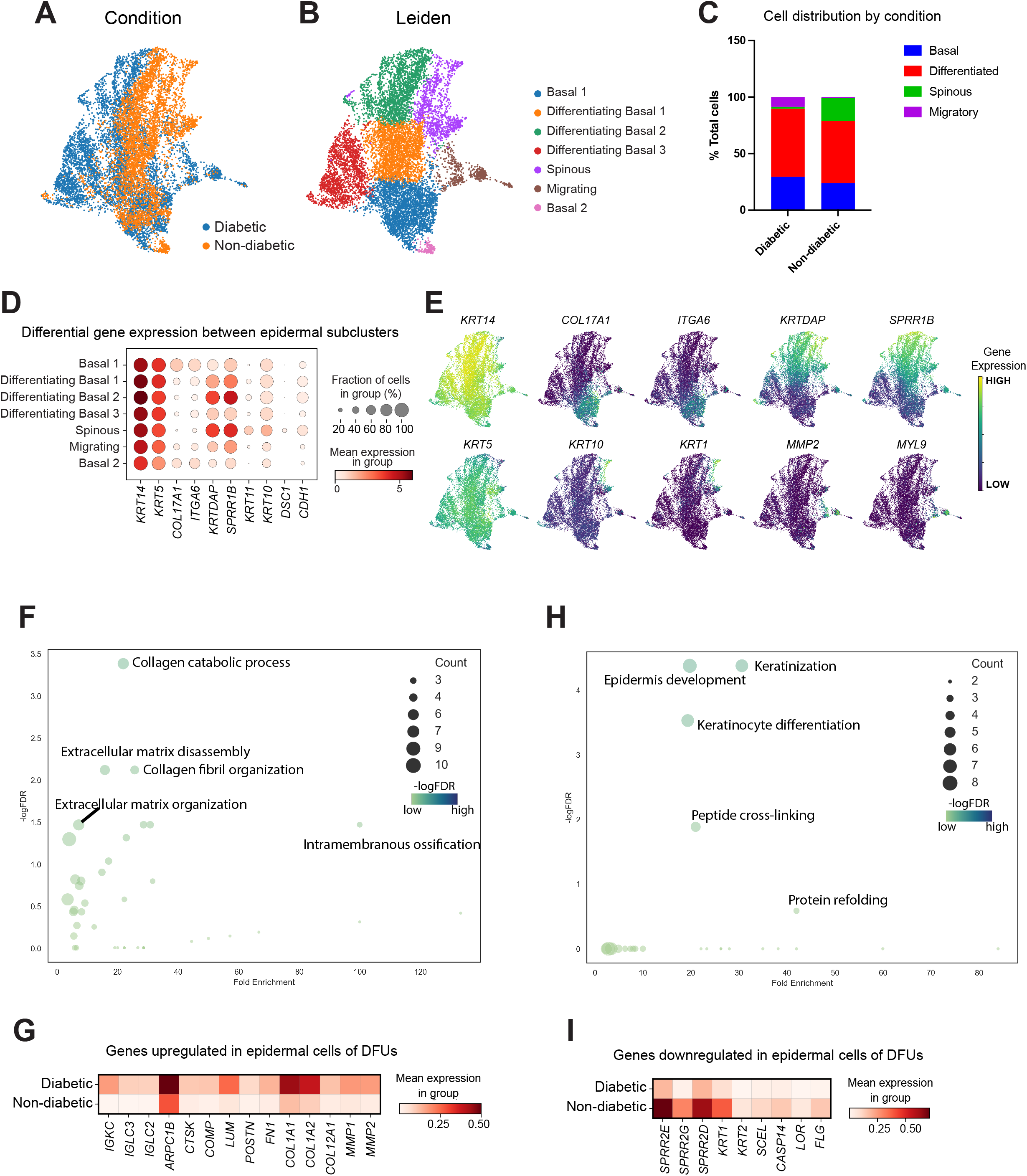
Analysis of epidermal cells reveals differences in epidermal sub-populations and in gene expression between DFU and NDFU. **(A)** UMAP clustering of epidermal cells by condition showing the distribution of DFU and NDFU cells in clusters. **(B)** UMAP clustering of epidermal cells showing the presence of epidermal subtypes. Clusters are identified by color and cell type. **(C)** Stacked bar graph showing the percentage of major subgroups of epidermal cells by condition. (n.s., not significant). **(D)** Dot plot showing the differential gene expression of markers used to identify each epidermal cell sub-cluster. The area of the circles indicates the proportion of cells expressing the gene, and the color intensity reflects the expression intensity. **(E)** Feature plots showing the expression of markers used to identify cell sub-clusters. The color intensity reflects the expression intensity. **(F)** GO analysis of biological processes downregulated in epidermal cells of DFU by -logFDR and Fold Enrichment. The area of the circles indicates the gene count, and the color intensity reflects the -logFDR power. **(G)** Matrix plot showing the differential expression of selected genes that are downregulated in epidermal cells of DFU. The color intensity reflects the expression intensity. **(H)** GO analysis of biological processes upregulated in epidermal cells of DFU by -logFDR and Fold Enrichment. The area of the circles indicates the gene count, and the color intensity reflects the -logFDR power. **(I)** Matrix plot showing the differential expression of selected genes that are upregulated in epidermal cells of DFU. The color intensity reflects the expression intensity. DFU, diabetic foot ulcers. FDR, false discovery rate. NDFU, non-diabetic foot ulcers. UMAP, uniform manifold approximation and projection.

To characterize the transcriptional changes in keratinocytes between diabetic and nondiabetic foot wounds, we analyzed the gene ontology of mRNAs altered in keratinocytes of diabetic patients (**Figure 2F-2I, Table 3**). Keratinocytes in diabetic foot wounds downregulated genes involved in keratinocyte differentiation including *SPRR2E, SPRR2G, KRT1, KRT2, SCEL, CASP14, LOR*, and *FLG* (**Figures 2F, 2G**). Keratinocytes in diabetic foot wounds increased genes involved in proteolysis and ECM organization/disassembly including *IGKC, IGLC2-3, ARPC1B, CTSK, COMP, LUM, POSTN, FN1, COL1A1, COL1A2, COL12A1, MMP1*, and *MMP2* (**Figures 2H, 2I**). This analysis also indicates that alterations in keratinocyte transcriptional profiles drive the major mRNA expression changes between control and diabetic foot wound samples (**Figure 1F-I**).

### Fibroblast lineages are altered in foot wounds of diabetic patients

Next, we analyzed changes in diabetic foot wound fibroblasts compared to control fibroblasts. Fibroblasts from all samples clustered into 5 groups with 3 unique clusters of fibroblasts in the diabetic samples (**Figure 3B-3C, S2A, Table 4)**. One cluster expressed microfibrillar-associated protein 5 (*MFAP5*) (**Figure 3B-3D**), which is secreted by cancer-associated fibroblasts in the mammary stroma and induces the invasion and migration of mammary tumor cells (Chen et al., 2020). A second cluster expressed *BMP4*, which has differential impact on myofibroblast formation in different tissues, including inhibiting fibrosis in the lung (Guan et al., 2022), and promote ossification in fibrodysplasia ossificans progression (Shafritz et al., 1996). Two clusters were enriched for *IL-6:* the first one, we named Inflammatory 1, was found in DFUs, and the second one, that we named Inflammatory 2, contained non-overlapping DFU and NDFU fibroblasts (**Figure 3A-3D)**. The major non-diabetic fibroblast cluster expressed the retinoic acid binding protein *CRABP1*, which is similar to fibroblasts in large mouse wounds and is expressed by fibroblasts in the upper dermis (Guerrero-Juarez et al., 2019) (**Figure 3A-3D**). Gene ontology analysis of mRNAs significantly altered in DFU fibroblasts compared to NDFU fibroblasts revealed that DFUs upregulated genes associated with ECM organization and cell adhesion (**Figure 3E, Table 3**). Consistent with altered ECM gene expression, analysis of genes associated with fibrosis, revealed that the *MFAP5* and *BMP4* clusters in the DFU samples and the *CRABP1* cluster in NDFU were highly enriched in fibrotic genes (**Figure S2B**).

**Figure 3.**
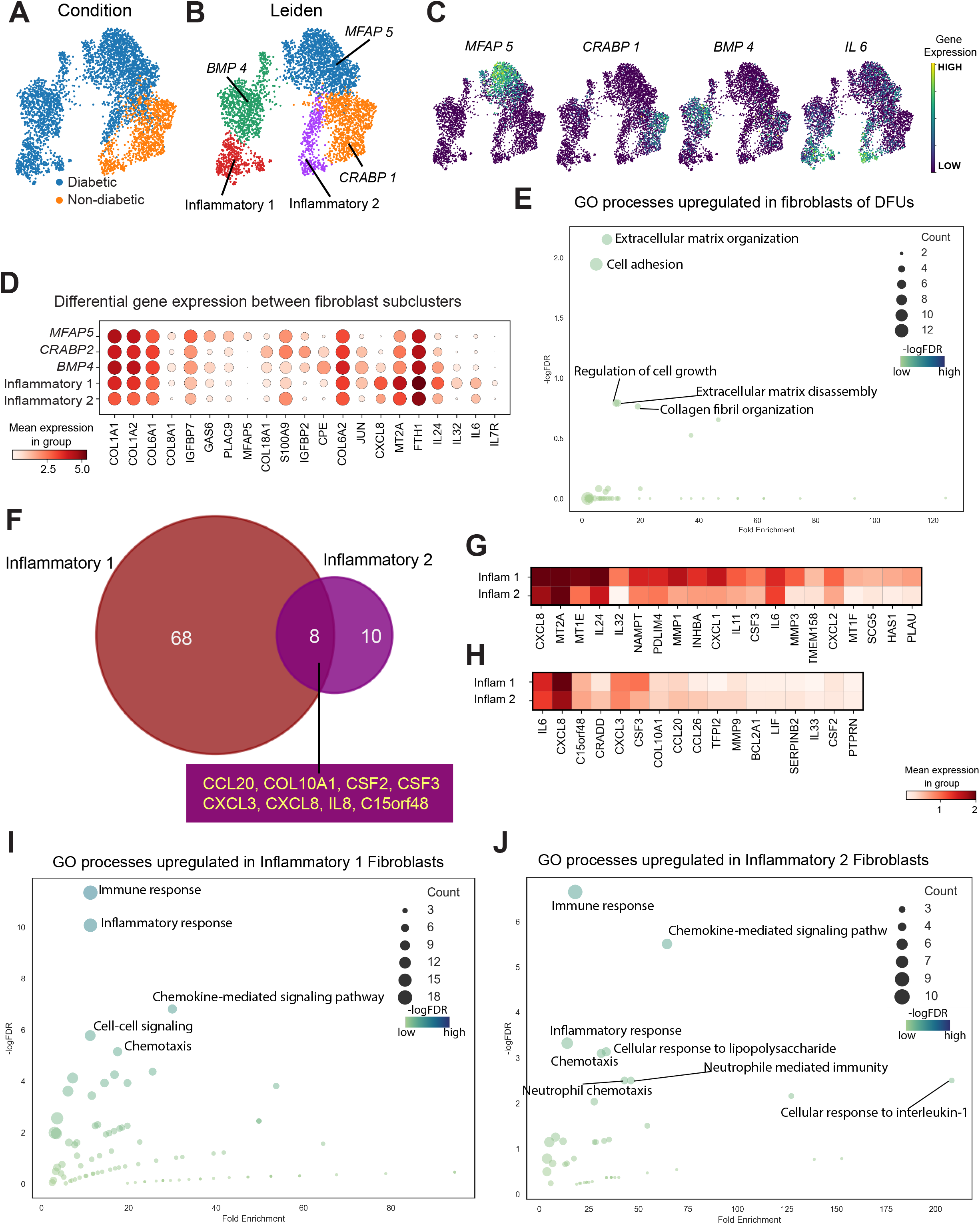
Analysis of Dermal Fibroblasts reveals differences in fibroblast sub-populations and in gene expression between DFU and NDFU. **(A)** UMAP clustering of dermal fibroblasts (DF) by condition showing the distribution of DFU and NDFU cells by cluster. **(B)** UMAP clustering of DF showing the presence of fibroblast subpopulations. Clusters are identified by color and either genetic marker (*MFAP1, CRABP1, BMP4*) or designated name (inflammatory 1, inflammatory 2). **(C)** Feature plots showing expression of markers used to identify DF subclusters. The color intensity reflects the expression intensity. **(D)** Dot plot showing the differential gene expression of markers used to identify each DF sub-cluster. The area of the circles indicates the proportion of cells expressing the gene, and the color intensity reflects the expression intensity. **(E)** GO analysis of biological processes upregulated in DF of DFU by - logFDR and Fold Enrichment. The area of the circles indicates the gene count, and the color intensity reflects the -logFDR power. **(F)** Trajectory and pseudo time analysis revealing the distinction between DFU specific fibroblasts (*BMP4* and Inflammatory 1) and non-DFU fibroblasts. **(G, H)** Matrix plot showing the differential expression of selected genes that are upregulated in inflammatory 1 (DFU) and inflammatory 2 (NDFU) fibroblasts, respectively. The color intensity reflects the expression intensity. **(I, J)** GO analysis of biological processes upregulated in inflammatory 1 and inflammatory 2 fibroblasts, respectively, by -logFDR and Fold Enrichment. The area of the circles indicates the gene count, and the color intensity reflects the -logFDR power. DF, dermal fibroblasts. DFU, diabetic foot ulcers. FDR, false discovery rate. NDFU, non-diabetic foot ulcers. PAGA, Partition-based graph abstraction. UMAP, uniform manifold approximation and projection.

We hypothesized that the fibroblast clusters could reveal altered developmental trajectories in the diabetic milieu. Thus, we sought to reconstruct the transitions that led to distinct clusters of fibroblasts in DFUs. To do so, we used Partition-based graph abstraction (PAGA) to reconcile Leiden clustering with trajectory inference to predict the relationships among the distinct fibroblast clusters. This analysis revealed two groups of fibroblast clusters that were not connected to each other. NDFU clusters CRABP1, Inflammatory 2, and DFU cluster MFAP5 were associated with each other, while DFU clusters BMP4 and Inflammatory 1 were connected to each other and separated from the rest of the clusters (**Figure 3F**). These connections were further confirmed by diffusion pseudotime analysis which estimates the developmental trajectories from a common “root cell”.

We individually assigned each of the fibroblast clusters CRABP1, MFAP5, and BMP4 as the root cells for the diffusion pseudotime analysis to investigate the relationship within the clusters. We found that the three associated clusters MFAP5, CRABP1, and Inflammatory 2 belonged to a common fibroblast lineage, while the DFU clusters BMP4 and Inflammatory 1 seemed to arise from a distinct developmental trajectory (**Figure S2D**).

We wanted to further characterize the two populations of fibroblasts that were enriched for *IL-6* expression, especially since our prior work identified a population of immune fibroblasts in non-healing DFUs compared to healing DFUs (Theocharidis et al., 2022). Thus, we compared the transcriptional profiles of the DFU clusters Inflammatory 1 and NDFU Inflammatory 2 (**Figure S2C**). The DFU Inflammatory 1 cluster upregulated several cytokines such as *IL24, CXCL8*, and several *MMPs* compared to the Inflammatory 2 cluster (**Figures 3G, 3I)**, which is like the inflammatory fibroblasts that we identified in a nonhealing DFUs previously (Theocharidis et al., 2022). By contrast, Inflammatory 2 upregulated genes involved in the cellular response to lipopolysaccharide, *TNF*, and *IL-1* as well as chemotaxis of neutrophils, monocytes, and leukocytes (**FIgures 3H, 3J**).

Another fibroblast-like cell, pericytes, also clustered separately from the main fibroblast populations (**Figure 1B, S1B**). DFUs upregulated genes involved in chemotaxis of immune cells and inflammation and downregulated genes involved in melanocyte biology (**Figure S3B-C**). Specifically, mRNAs for different signaling molecules in the BMP, WNT, and interleukin family were upregulated in pericytes of DFUs compared to NDFUs (**Figure S3D**).

### Macrophages alter ECM and inflammation genes in foot wounds of diabetic patients

Given the differences in inflammatory fibroblasts in NDFU and DFU, we clustered CD45+ immune cells alone. While mast cells, macrophages, and T cells were present in both DFUs and NDFUs, plasma cells seem to be enriched in DFUs only (**Figure 4A-E**). Each cell type showed some transcriptional differences between NDFU and DFU (**Figure 4A**), yet the major gene expression changes were only found in macrophages, which have been implicated in DFU pathology (Joshi et al., 2020, Mirza and Koh, 2011).

**Figure 4.**
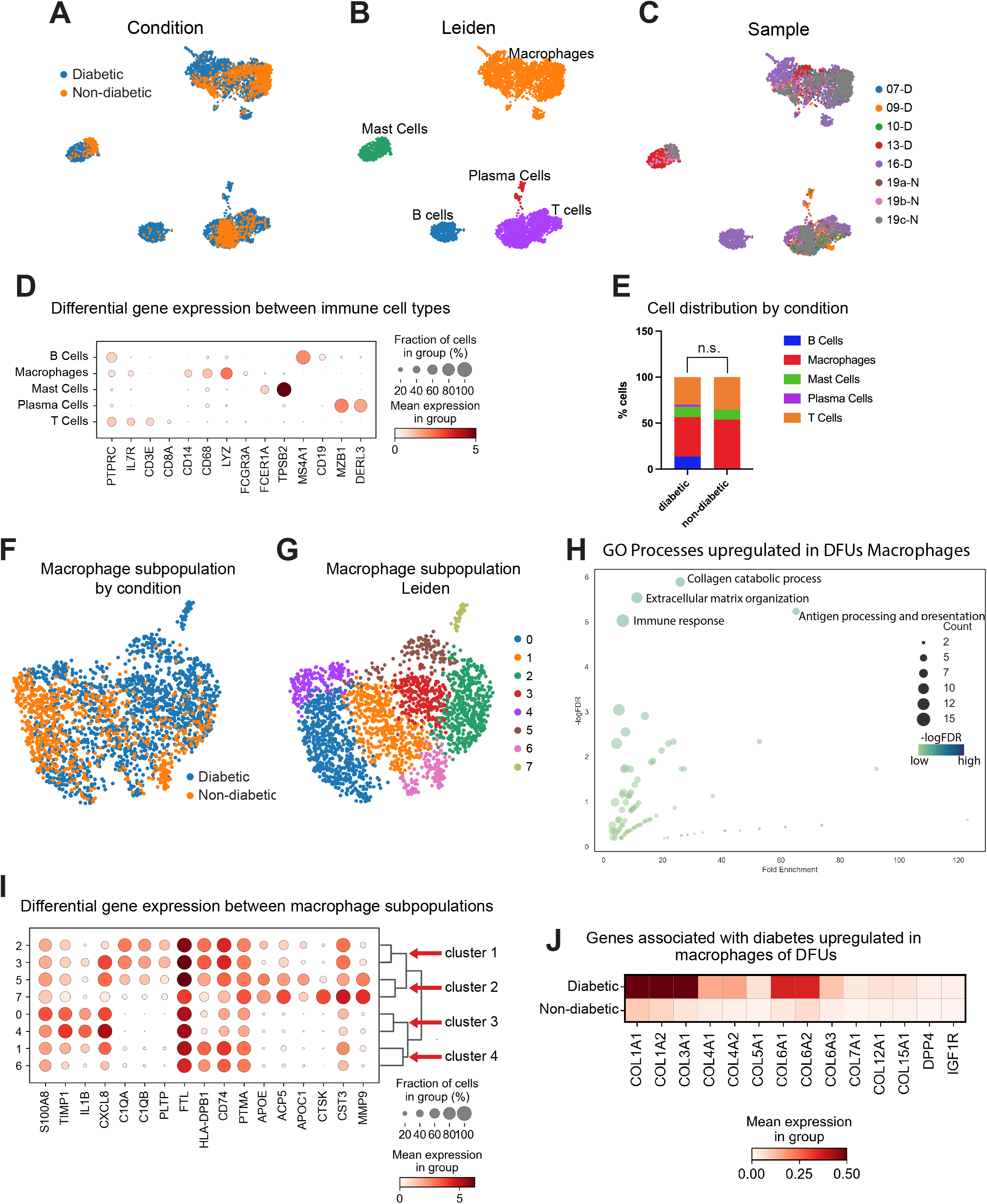
Analysis of immune cells reveals differences in macrophage sub-populations and gene expression between DFU and NDFU. **(A)** UMAP clustering of immune cells by condition showing the distribution of DFU and NDFU cells in clusters. **(B)** UMAP clustering of immune cells subpopulations. Clusters are identified by color and cell type. **(C)** UMAP clustering of individual samples showing the contribution of each sample to the clusters. **(D)** Dot plot showing the differential gene expression of markers used to identify each immune cell subpopulation. The area of the circles indicates the proportion of cells expressing the gene, and the color intensity reflects the expression intensity. **(E)** Stacked bar graph showing the percentage of immune cells subpopulations by condition. (n.s., not significant). **(F)** UMAP clustering of macrophages by condition showing the distribution of DFU and NDFU cells in clusters. **(G)** UMAP clustering of macrophage subpopulations. Clusters are identified by color and number. **(H)** GO analysis of biological processes upregulated in macrophages of DFU by - logFDR and Fold Enrichment. The area of the circles indicates the gene count, and the color intensity reflects the -logFDR power. **(I)** Dot plot showing the differential gene expression of markers used to identify each macrophage subpopulation. Given the similarities in gene expression, macrophage clusters have been grouped in 4 sub-clusters. The area of the circles indicates the proportion of cells expressing the gene, and the color intensity reflects the expression intensity. **(J)** Matrix plot showing the differential expression of genes associated with diabetes that are upregulated in macrophages of DFU. The color intensity reflects the expression intensity. DFU, diabetic foot ulcers. FDR, false discovery rate. NDFU, non-diabetic foot ulcers. UMAP, uniform manifold approximation and projection.

Unbiased clustering of specific macrophage clusters from healthy and diabetic foot wounds revealed distinct clusters of macrophages in NDFU and DFU (**Figure 4F, 4G**). To understand these differences, we interrogated gene expression changes between macrophages in DFUs and NDFUs. Macrophages from DFUs significantly upregulated mRNAs involved in ECM organization and immune response (**Figure 4H, Table 3)**. Consistent with these categories, we find that macrophages upregulate many ECM genes involved in fibrosis (**Figure 4I**).

Specific analysis of differentially expressed genes in the 7 macrophage clusters revealed subclusters of pairs of macrophage clusters. Subclusters 1 and 2 were more prevalent in DFUs and expressed mRNAs encoding for complement, while subclusters 3 and 4 were mixed between DFUs and NDFUs (**Figure 4F-I**). Moreover, macrophages of DFUs overexpressed several collagen genes, and genes involved in insulin signaling and metabolism, like *DPP4* and *IGFR1* (**Figure 4J**), all of which have been associated with diabetes (Fernández et al., 2001, Liu et al., 2022, Röhrborn et al., 2015).

Endothelial cells also displayed differences in DFUs compared to NDFUs (**Figure S4A-S4C**). DFU endothelial cells upregulated genes involved in ECM and immune regulation (**Figure S4D, Table 3**) and downregulated genes involved in humoral response and autocrine signaling (**Figure S4E**).

### Predicted cellular communication of stromal cells is altered in foot wounds of diabetic patients

Our prior work and that of others have shown that fibroblasts and macrophages communicate in tissues during inflammation and repair (Buechler et al., 2021a, Shook et al., 2018, Zhou et al., 2018). To interrogate the cellular interactions of stromal cells in more depth, we utilized the CellChat algorithm (Jin et al., 2021), which infers intercellular communication networks from scRNA sequencing data. We noted that the number and strength of the predicted intercellular interactions differed between non-diabetic and diabetic samples (**Figure 5A**). In particular, new interactions were predicted between macrophages and T cells, mast cells, and fibroblasts in DFUs compared to NDFUs (**Figure 5A**). Furthermore, the strength of interactions of endothelial cells, fibroblasts, pericytes, and mast cells with T cells were predicted to be elevated in DFUs, while the inferred interactions of DFU lymphatic vessels were reduced with T cells (**Figure 5A**).

**Figure 5.**
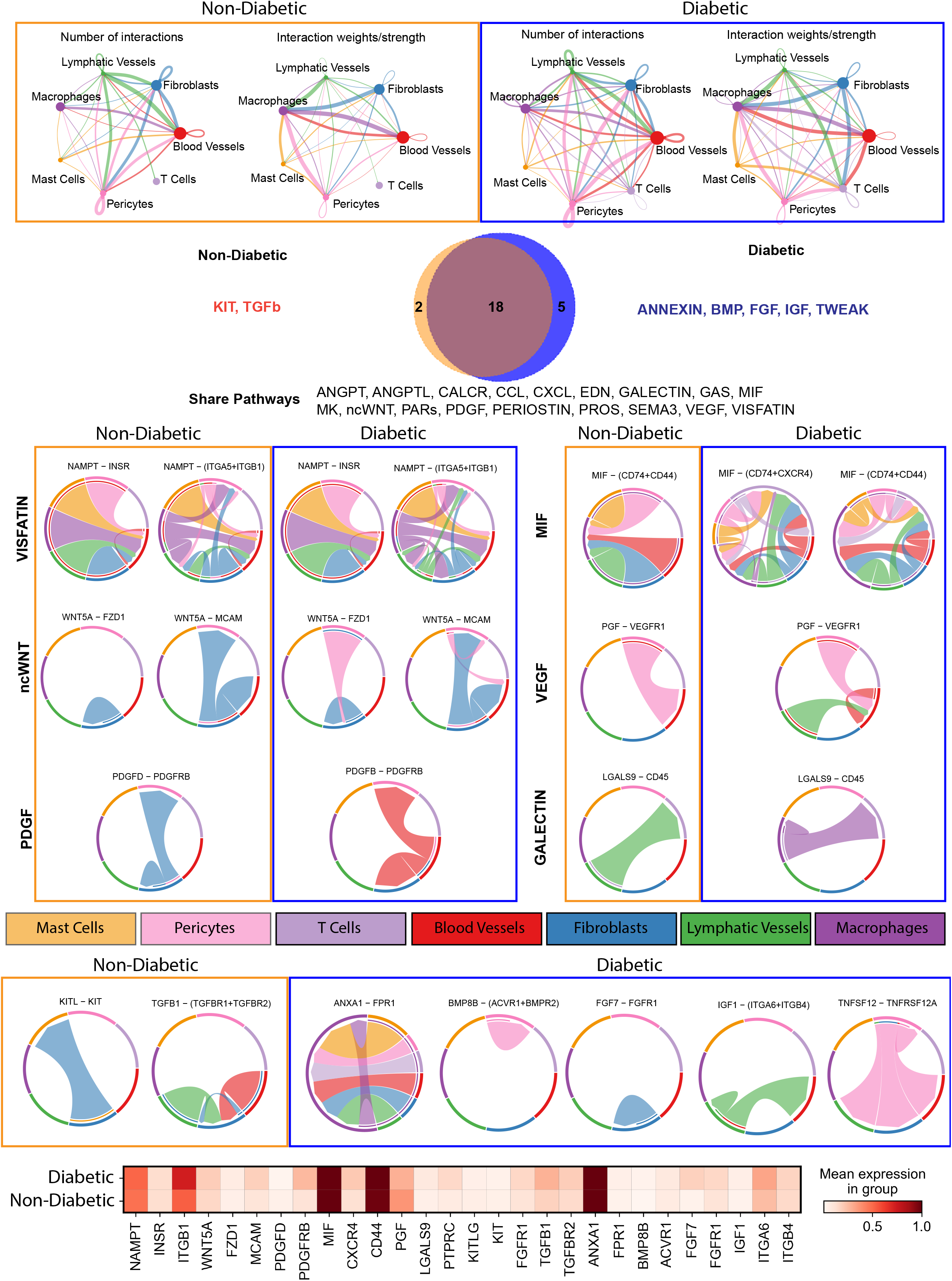
Cell-cell communication analysis of multiple cell types comparing DFU to NDFU samples. **(A)** Circle plots showing the aggregated cell-cell communication networks (number of interactions and interactions strength) of non-DFU and DFU samples. **(B)** Venn diagram showing the overlapping and distinct significant pathways of each condition. **(C)** Chord diagrams showing the significant pathways between non-DFU and DFU samples related to wound healing: *VISFATIN, ncWNT, PDGF, MIF, VEGF, GALECTIN*. The specific cell types are color-coded as indicated in the bars. **(D)** Chord diagrams showing the specific pathways of non-DFU samples: *KIT, TGFb*. **(E)** Chord diagrams showing the specific pathways of DFU samples: *ANNEXIN, BMP, FGF, IGF, TWEAK*. **(F)** Matrix plot showing the expression levels of ligands and receptors from each significant pathway. DFU, diabetic foot ulcers. NDFU, non-diabetic foot ulcers. UMAP, uniform manifold approximation and projection.

CellChat predicted interactions of 18 specific signaling pathways in both non-diabetic and diabetic foot wounds (**Figure 5B**). Visfatin/nicotinamide phosphoribosyltransferase (NAMPT), a proinflammatory cytokine associated with insulin resistance (Dahl et al., 2012) was predicted to interact with the insulin receptor (INSR) on blood vessels similar in both conditions, and only slight differences in interactions with the integrins on several cell types (**Figure 5C**). However, other predicted pathways display altered interactions between cells in DFUs compared to NDFUs. For instance, both non-canonical Wingless (ncWNT) signaling via Wnt5A and placental growth factor (PGF) displayed additional interactions in DFUs. In NDFUs, pericytes were predicted to produce WNT5A to activate Frizzled (FZD)1 receptors on fibroblasts in DFUs but not NDFUs (**Figure 5C**). For PGF, vascular endothelial growth factor receptor (VEGFR) on blood endothelial cells in DFUs displayed weaker interactions between PGF-derived from pericytes and new interactions with PGF-derived from autocrine signaling and fibroblast derived-PGF.

CellChat also predicted that DFUs altered ligand sources and types compared to NDFUs. Activation of Platelet derived growth factor receptor (PDGFR)B on pericytes and fibroblasts via fibroblast derived PDGFD was inferred in NDFUs, while only PDGFB from blood endothelial cells was predicted in DFUs (**Figure 5C**). Interestingly, dysregulation of the galectin-9 (LGALS9) pathway that is emerging as an indicator of disease severity (Moar and Tandon, 2021) and associated with diabetes (Sun et al., 2020), was predicted to activate CD45 on T cells by lymphatic vessel-derived LGALS9 in NDFUs, whereas LGALS9 was inferred to be macrophage derived in DFUs **(Figure 5C**). The most dramatic changes in interactions were predicted for macrophage migration inhibitory factor (MIF) signaling, a cytokine that contributes to immunity and tissue homeostasis (Jankauskas et al., 2019). CD74+CD44 on macrophages was predicted to be activated by MIF from pericytes, blood and lymphatic vessels, and fibroblasts, whereas DFUs were predicted to also activate T cells from MIF derived from these cell types as well as mast cells. Furthermore, MIF from all stromal cells analyzed were predicted to produce MIF to also activate CD74+CXCR4 on fibroblasts and T cells **(Figure 5C**).

In addition to alterations in shared pathways, DFUs lacked signaling via c-kit ligand (KIT) and (Transforming Growth Factor) TGFβ, which were inferred to be active in NDFUs. TGFβRs on NDFUs fibroblasts were predicted to be activated by autocrine and paracrine lymphatic and blood endothelial-derived TGFβ signals (**Figure 5D**). For KIT, fibroblasts in NDFUs expressed KIT, which was predicted to activate KITR on mast cells in non-diabetic samples (**Figure 5D**).

Five pathways were significantly predicted in diabetic samples and not non-diabetic wounds. Anxa1 ligands derived from all stromal cells and autocrine signals were inferred to activate formyl peptide receptor 1 (FPR1) on macrophages, which may increase polarization toward anti-inflammatory cells (McArthur et al., 2020) (**Figure 5E**). Pericytes in DFUs also upregulated *BMP8B* and was predicted to initiate autocrine signaling via BMPR2 (**Figure 5E**), and TNFSF12 was predicted to activate autocrine and paracrine signaling to TNFRSF12A on fibroblasts and endothelial cells. Finally, *IGF1* expression by lymphatic vessels was predicted to induce signaling via integrins A6 and B4 in autocrine and paracrine signaling on blood vessels in DFUs. Analysis of gene expression for these ligands and receptors displayed slight changes between DFUs and NDFUs, suggesting that cell type specific changes are likely to be responsible for these predicted changes in cell signaling (**Figure 5F**).

## Discussion

Although diabetic wounds have been analyzed in mice and humans at the single-cell level (Theocharidis et al., 2022), key mechanisms leading to the poor clinical outcome of these wounds are still not well understood. The proper and temporal coordination of multiple cell types is required for tissue repair, and the mechanisms that impact chronic wounds of diabetic foot skin remain unclear. Here, we provide a single-cell atlas of foot wounds in both non-diabetic and diabetic patients and investigate the key regulatory pathways and cellular heterogeneity of cell types in these wounds. These results will aid in understanding wound repair in volar skin and diabetic healing in depth and provide targets for clinical therapies of these diseases.

Many studies have indicated several defects in diabetic wounds including keratinocyte proliferation, migration, and expression of inflammatory cytokines and ECM modifying enzymes, like MMPs (Hu and Lan, 2016, Stojadinovic et al., 2008). While keratinocyte re-epithelialization migration and proliferation is important for wound repair (Eming et al., 2014), our data suggest that a major defect in DFUs is an impairment in differentiation in which keratinocytes reform the stratified epithelium to restore the essential barrier of the epidermis, which is consistent with other studies noting a maintenance of keratinocytes in a proliferative and reduced differentiated state in chronic wounds (Andriessen et al., 1995, Hu and Lan, 2016, Stojadinovic et al., 2008). Interestingly, differentiation defects also are hallmarks of other inflammatory skin diseases like atopic dermatitis and psoriasis (Wikramanayake et al., 2014). Perhaps the upregulation of inflammatory factors by fibroblasts and macrophages in diabetic keratinocytes identified in our study could block keratinocyte differentiation (Wolf et al., 2021)’(Boniface et al., 2005). Defects in ECM production and remodeling by stromal cells could also alter DFU keratinocyte differentiation, which requires loss of integrin-based adhesion to the underlying basement membrane and dermis (Li et al., 2007, Martin, 1997).

Fibroblasts are another central regulator of skin repair, driving repair of dermal ECM and regulation of the inflammatory response (Plikus et al., 2021). Here, we find that DFU fibroblasts broadly induced genes that control ECM organization and inflammation compared to NDFUs (Figure 3). Importantly, NDFUs contained a population of CFAP5^hi^ high fibroblasts that are associated with regenerative repair in mice and reindeer (Guerrero-Juarez et al., 2019, Sinha et al., 2022), and these populations were absent from the DFUs. Instead, our data suggests that diabetes induces distinct fibroblast transitions from a CRABP1^hi^ population to fibroblasts that express different ECM regulators including MFAP5, which is also found in tumor-associated fibroblasts and promotes cancer cell invasion and migration (Chen et al., 2020). Furthermore, we identified unique inflammatory fibroblasts in DFUs, which is consistent with our prior work that showing an inflammatory fibroblast population in healing DFUs but not nonhealing DFUs (Theocharidis et al., 2022). Interestingly, we find that DFUs harbor fibroblast populations that transcriptionally precede the DFU-specific inflammatory population suggesting that diabetes leads to generation of unique fibroblasts in foot wounds. These fibroblast populations here could not be classified into previously-defined fibroblast subclusters based on *SFRP1, SFRP2*, and *CXCL12* axes in non-diabetic human skin (Ascensión et al., 2021) or the proposed universal fibroblast markers in mice and human steady-state or inflamed state (Buechler et al., 2021b)(**Figure 3C, 3D and S2A**). That said, BMP4+ fibroblasts were noted in the mouse intestine (Buechler et al., 2021b). Thus, our data point to a tissue and disease state specific influence on fibroblast phenotype.

Chronic, non-healing DFUs harbor persistent inflammatory macrophages in both diabetic human samples (Theocharidis et al., 2022) and mouse models (Joshi et al., 2020, Mirza and Koh, 2011). We found that DFU macrophages highly altered genes involved in antigen presentation and ECM remodeling and developed subpopulations unique to diabetic wounds. One diabetic specific macrophage population upregulated mRNAs encoding genes involved in the complement system, which has been shown to be elevated in DFUs (Kheiralla et al., 2012), and the other induced genes that regulate macrophage polarization and ECM degradation. Interestingly, the LGALS9 signaling pathway that is aberrant in DFUs including a unique autocrine activation in macrophages that is absent in NDFUs, may contribute to polarization defects since LGALS9 signaling has been implicated in macrophage polarization (Zhang et al., 2019). Thus, macrophages likely contribute to ECM and inflammatory alterations in DFUs.

Given these differences in gene expression and cell populations, many cellular interactions were predicted to be perturbed including more and stronger interactions between cell types including T cells, which have been shown to have lower differentiation in DFUs (Moura et al., 2017). Furthermore, we identify several pathways that may be upregulated to compensate for aberrant repair in DFUs. We found that DFUs displayed induced ANXA1 signaling to macrophages, which has also shown to facilitate wound closure in genetically diabetic mice (Huang et al., 2020) and improve recovery from diabetic nephropathy in mice (Wu et al., 2021). Other factors induced in DFUs include (Tumor necrosis factor weak inducer of apoptosis) TWEAK signaling, which promotes healing in human burn wounds (Liu et al., 2018) and is associated with inflammation in obesity (Vendrell and Chacón, 2013), and FGF7/KGF, which is required for healing in skin wounds (Peng et al., 2011). It is possible, given that these pathways can promote healing, that DFUs may induce these pathways to compensate for defects in healing. Alternatively, defects in timing of these pathways in the wounding process may promote the defects in DFUs. Future studies defining the impact of these pathways on DFUs are needed to fully understand their impact.

While the pathways specific to DFUs may be compensatory, the absence of KIT and TGFb signaling in DFUs could contribute to defects in healing. Kit signaling promotes myocardial healing after infarction (Cimini et al., 2007), corneal wound repair (Miyamoto et al., 2012), and may improve healing in skin wounds (Zgheib et al., 2015). Furthermore, TGFb signaling contributes to tissue repair in a context and isoform specific manner (Divoux et al., 2010), and may impair fibroblast ECM regulation, driving fibrosis (Shi-wen et al., 2010) or generating aberrant fibroblast populations identified here. Taken together, our study provides evidence for aberrant transcriptional changes, cellular development, and communication in DFUs that could provide a platform for improving chronic diabetic injuries and importantly, the mechanisms that might drive human volar skin repair.

## Methods

### Subjects

Diabetic and non-diabetic adults with chronic foot ulcers that were undergoing skin wound debridement were consented to donate discarded tissue for this study (IRB approval # 1609018360)(**Table 1**). The diabetic foot ulcer specimens were obtained from 5 individuals diagnosed with Type 2 diabetes. The non-diabetic foot ulcer specimens were obtained from the same individual, 2 of which were from the same wound, one month apart from each other.

### Tissue collection and processing

Skin wound specimens were collected at the clinical setting in PBS with 1% Pen/Strep and transported to the lab on ice for processing. All specimens were processed within 3 hr. of collection. The specimen was cleaned by sequentially immersing it in 10% Betadine, twice in 70% ethanol, and twice in PBS for 1-min at a time. Tissue was then placed on a petri dish to remove excess blood and subcutaneous fat, sliced into 0.5cm pieces and incubated in 5U/ml dispase (Stemcell Technologies, 07913) overnight at 4°C. The next day, the epidermis was carefully peeled off with tweezers and both the epidermis and dermis were finely minced with a No. 10 sterile scalpel and scissors. The minced tissue was then placed in an enzyme cocktail consisting of 1U/ml dispase (Stemcell Technologies, 07913), 2,000U/ml collagenase I (Worthington, M3A14008A), and 424U/ml collagenase II (Worthington, LS004176) in 0.25% Trypsin-EDTA (Gibco, 25200-056), at 37 °C for 1 hr. with constant shaking. Enzymes were inactivated by adding an equal volume of complete DMEM (+10% FBS, +1%Pen/Strep). The cell suspension was then passed through 70mm and 40mm cell strainers and centrifuged for 10 min at 500 x *g* at 4 °C. For blood cell lysis, the cell pellet was incubated with ACK buffer (Lonza, 10-548E) for 5min. at RT. The lysis was stopped with complete DMEM (+10% FBS, +1%Pen/Strep) and centrifuged for 5min at 500 x *g*. The pellet was resuspended in 500 ml of complete DMEM (+10% FBS, +1%Pen/Strep) and passed through a 40mm cell strainer. The final single-cell solution was resuspended in DMEM with 0.04% BSA and adjusted to a concentration of 700-1200 cells/ml.

### 3’-end single cell gene expression

10X Genomics 3’-end single cell gene expression libraries were performed using Single Cell 3’ v2 chemistry (10X Genomics). Libraries were sequenced on Illumina to achieve reads depth per sample. Fastqs file from sequencing were aligned to GRCh38 reference genome using Cellranger version 3.0.2. The Cellranger output h5 files were used for downstream analysis. All samples were combined into a large anndata (version 0.7.5) object to be analyzed using Scanpy version 1.6.0. The samples were labeled with Age, Sex, and Condition (diabetic and non-diabetic).

### QC metrics

We filtered out cells that expressed less than 200 genes and genes that were expressed in less than 3 cells. In addition, to eliminate doublets and low-quality cells, we also excluded cells that expressed more than 5000 genes and had higher than 15% mitochondrial genes. There were 27998 cells from 8 samples (5 diabetic and 3 non-diabetic) remained after the filtering steps. The gene expressions of these cells were normalized and log transformed for downstream analysis.

### Batch correction

To integrate the heterogeneous samples, we performed batch-correction using Scanorama (Hie et al., 2019).

### Downstream analyses

Principal Component Analysis of integrated samples were performed and used for k Nearest Neighbor graph construction. Uniform Manifold Approximation and Projection dimensionality reduction with default value was done. Leiden community detection with low resolution was done to classify different cell types clusters. Scanpy’s differential expression analysis rank_genes_groups with Wilcoxon was used to identify distinct cell types from each cluster. For analysis of each cell type, we subset them using the annotated markers from the main object.

Partition-based graph abstraction (PAGA) analysis was used to reconstruct branching gene expression changes across different fibroblast subsets to establish the lineage relations among them. Diffusion pseudotime analysis was used in conjunction with PAGA graph to predict the developmental trajectory of distinct fibroblast clusters (Haghverdi et al., 2015, Wolf et al., 2019).

Cell-cell communication analysis was done using CellChat version 1.6.0 (Jin et al., 2021).

### GO analysis

DEGs from each analysis were used for GO terms analysis using Database for Annotation, Visualization and Integrated Discovery (DAVID). The results were exported as csv file for visualization and downstream analysis.

## Data availability

Data sets related to this article can be found at https://skinregeneration.org/data/ and hosted at the Gene Expression Omnibus (GEO) (GSE223964).

## Acknowledgements

We would like to thank members of the Horsley laboratory for their feedback and critical analysis of these data and manuscript. This work was funded by the NIDDK-sponsored Diabetic Complications Consortium grant 5U24DK115255-04, and we value the feedback we received from the consortium, especially Drs. Aristidis Veves and Manoj Bhasin. V.H. is funded by N.I.H-NIAMS R01s AR076938, AR079232, and AR075412.

## Author Contributions

**Teresa Sandoval-Schaefer**: Investigation, Formal analysis, Visualization, and Writing – Original Draft; **Quan Phan**: Formal analysis, Conceptualization, Visualization, and Writing – Original Draft; **Biraja C. Dash**: Investigation; **Alexandre J. Prassinos**: Resources; **Kaiti Duan**: Resources; **Michael I. Gazes**: Resources; **Steven D. Vyce:** Resources; **Ryan Driskell:** Supervision, Writing – Original Draft; **Henry C. Hsia**: Resources, Conceptualization, Supervision, and Writing – Original Draft; **Valerie Horsley:** Conceptualization, Project Administration, Visualization, Funding Acquisition, and Writing – Original Draft.

## Inclusion and Diversity

One or more of the authors of this paper self-identifies as an underrepresented ethnic minority in their field of research or within their geographical location. One or more of the authors of this paper self-identifies as a gender minority in their field of research. While citing references scientifically relevant for this work, we also actively worked to promote gender balance in our reference list.

## Figure Legends

**Supplementary Figure S1.**
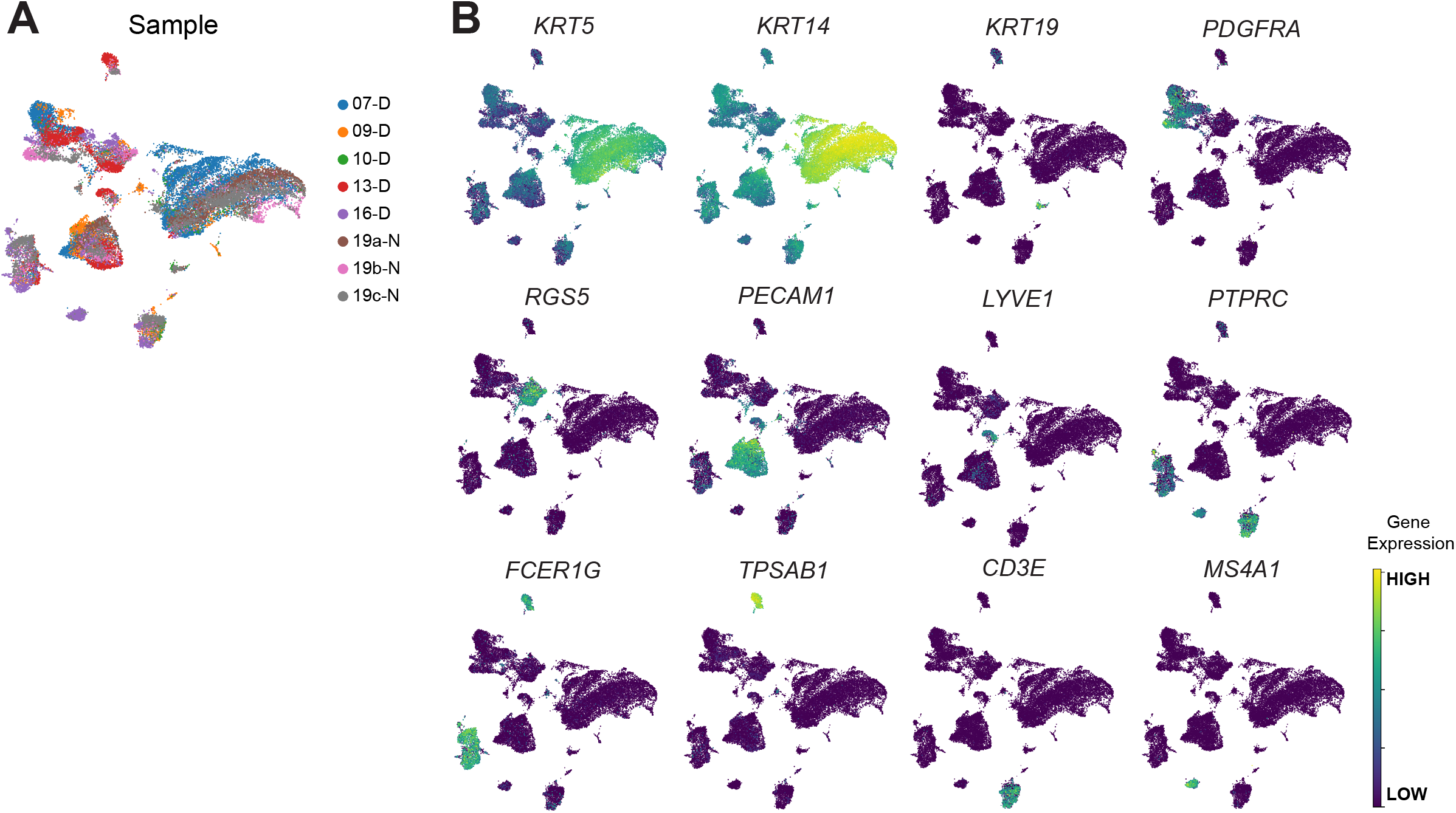
Contribution of individual samples to cell clusters and gene expression of specific markers. **(A)** UMAP clustering of individual samples showing the contribution of each sample to the clusters. **(B)** Feature plots showing expression of markers used to identify cell clusters. The color intensity reflects the expression intensity. UMAP, uniform manifold approximation and projection.

**Supplementary Figure S2.**
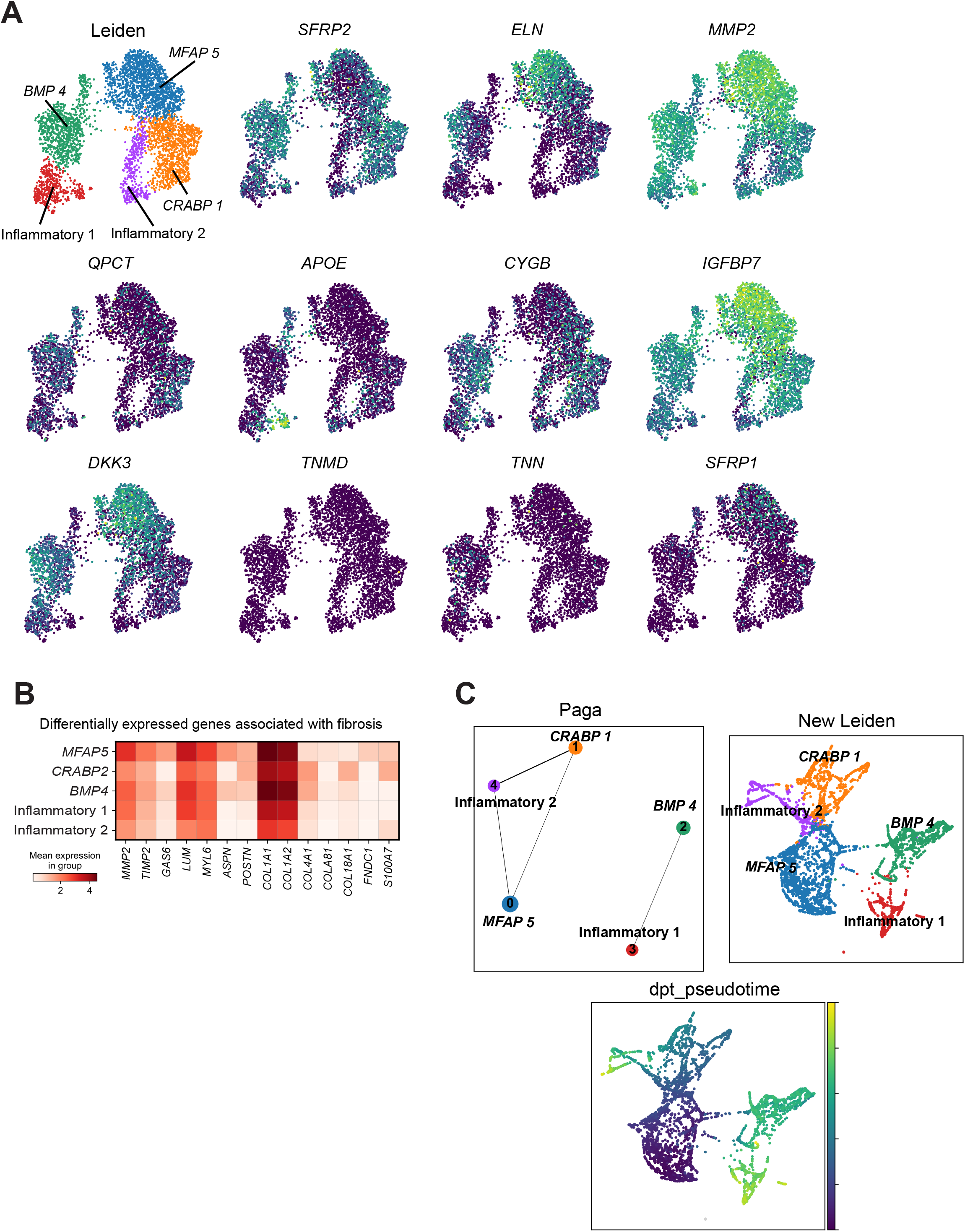
Subset analysis of dermal fibroblasts from DFU and NDFU samples. **(A)** Feature plots showing expression of known DF markers. The UMAP of DF subpopulations identified in our samples is provided for reference (top left). **(B)** Matrix plot showing the differential expression of genes associated with fibrosis across all five clusters. The color intensity reflects the expression intensity. **(C)** Venn diagram revealing the distinct and shared genes between the two inflammatory fibroblast clusters (highlighted genes are shared). **(D)** Pseudotime analysis depicting the *MFAP5, CRABP1*, or *BMP4* clusters as root of the inflammatory fibroblasts. DF, dermal fibroblasts. DFU, diabetic foot ulcers. NDFU, non-diabetic foot ulcers. PAGA, Partition-based graph abstraction. UMAP, uniform manifold approximation and projection.

**Supplementary Figure S3.**
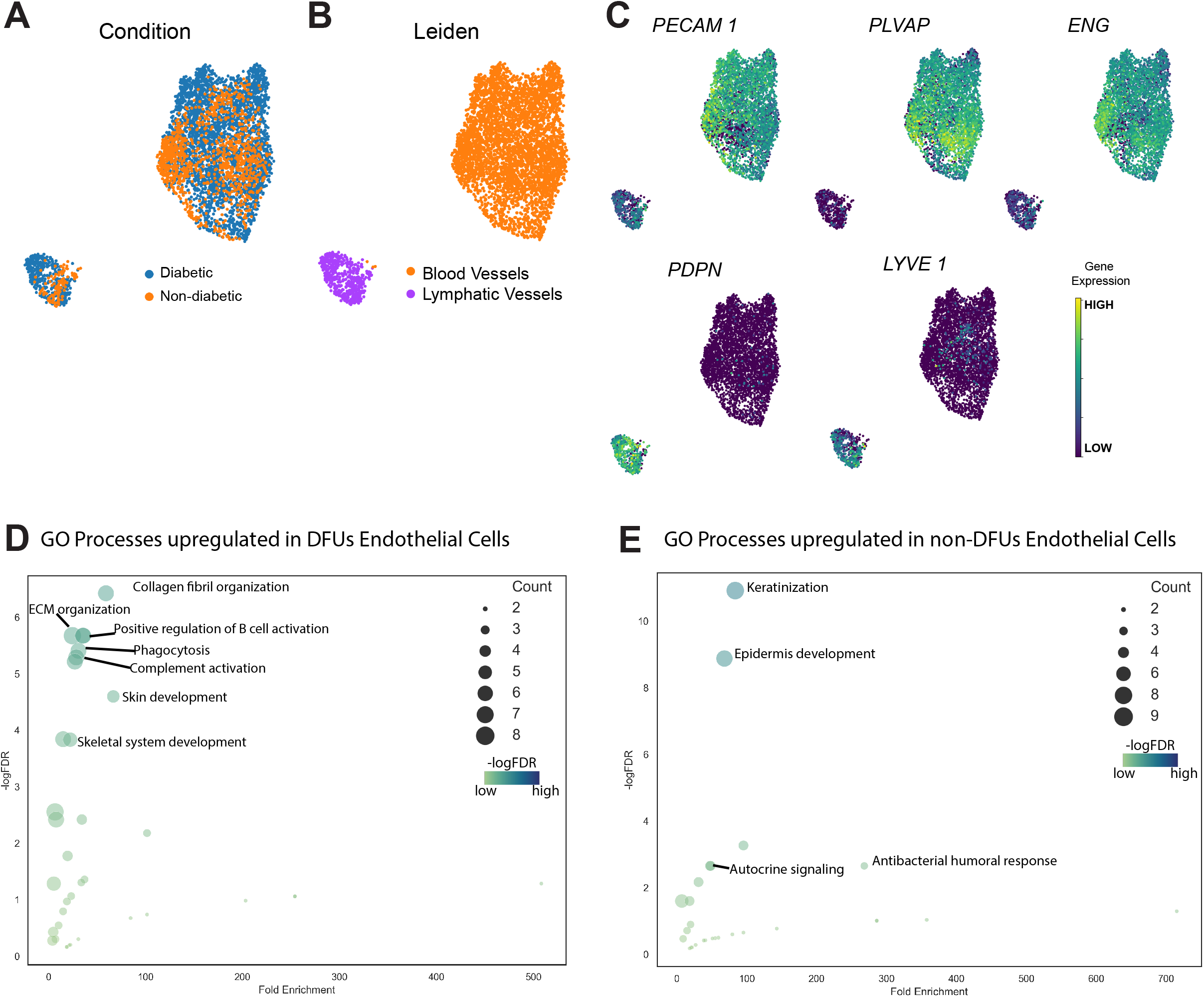
Analysis of pericytes reveals differences in gene expression between DFU and NDFU. **(A)** UMAP clustering of pericytes by condition showing the distribution of DFU and NDFU cells in clusters. **(B, C)** GO analysis of biological processes upregulated and downregulated, respectively, in pericytes of DFU by -logFDR and Fold Enrichment. The area of the circles indicates the gene count, and the color intensity reflects the -logFDR power. **(D)** Violin plots of selected genes involved in cell-cell signaling and immune response that are upregulated in pericytes of DFU. DFU, diabetic foot ulcers. FDR, false discovery rate. NDFU, non-diabetic foot ulcers. UMAP, uniform manifold approximation and projection.

**Supplementary Figure S4.**
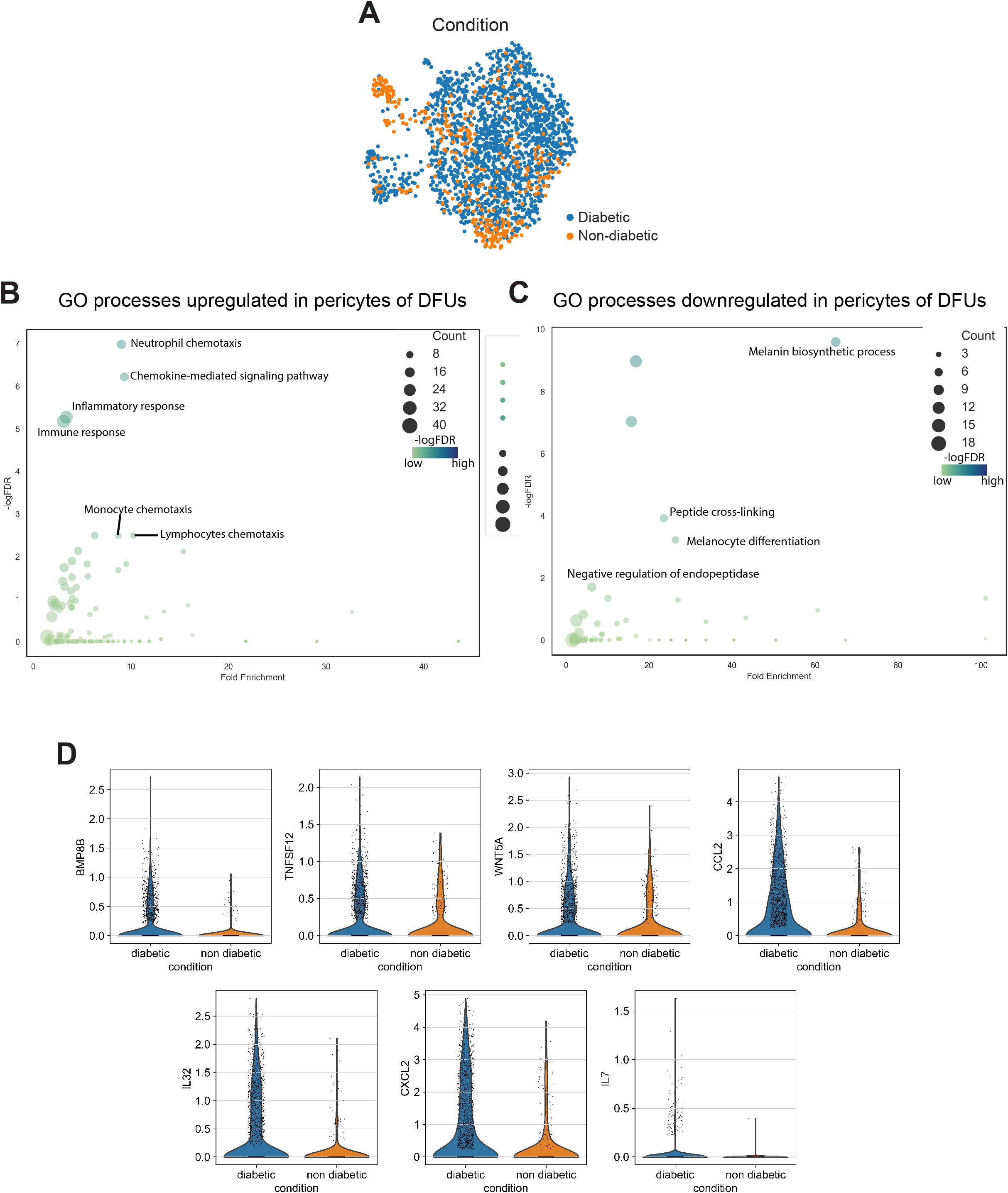
Analysis of endothelial cells reveals differences in gene expression between DFU and NDFU. **(A)** UMAP clustering of endothelial cells by condition showing the distribution of DFU and NDFU cells in clusters. **(B)** UMAP clustering of endothelial cells subtypes. Clusters are identified by color and cell type. **(C)** Feature plots showing the expression of markers used to discern blood vessels from lymphatic vessels. The color intensity reflects the expression intensity. **(D, E)** GO analysis of biological processes upregulated and downregulated, respectively, in endothelial cells of DFU by -logFDR and Fold Enrichment. The area of the circles indicates the gene count, and the color intensity reflects the -logFDR power. DFU, diabetic foot ulcers. FDR, false discovery rate. NDFU, non-diabetic foot ulcers. UMAP, uniform manifold approximation and projection.

