## Supplementary material for "Transcriptional heterogeneity in human diabetic foot wounds": Table 1

| Sample ID | Diabetic (Y/N) | Age at time of collection (years) | Gender | Race | Hispanic (Y/N) | Location of wound | Infection  (Y/N) | Epithelialization  (Y/N) | Size of wound on collection |
| --- | --- | --- | --- | --- | --- | --- | --- | --- | --- |
| 07-D | Y | 53 | F | Black | N | Plantar medial hallux | N | N | 0.04 sq cm |
| 09-D | Y | 63 | F | White | N | Chercot foot wound, plantar lateral | N | N | 0.91 sq cm |
| 10-D | Y | 54 | M | Black | N | Plantar forefoot wound sub metatarsals 3/4 | N | Y | 22 sq cm |
| 13-D | Y | 59 | M | White | Y | Foot plantar, first MPJ | N | Y | 1.2 sq cm |
| 16-D | Y | 67 | F | Other/Not listed | Y | Heel wound | N | Y | 24.4 sq cm |
| 19a-N | N | 49 | M | White | N | Left plantar midfoot wound | N | N | 14 sq cm |
| 19b-N | Same subject as above | -- | -- | -- | -- | Left plantar midfoot wound (follow-up, same wound as above) | N | N | 257 sq cm |
| 19c-N | Same subject as above | -- | -- | -- | -- | Right plantar midfoot wound | N | N | 17.5 sq cm |
